## Supplementary figures and images for "Mod3D: A Low-Cost, Flexible Modular System of Live-Cell Microscopy Chambers and Holders"

### Supplemental Figure 1

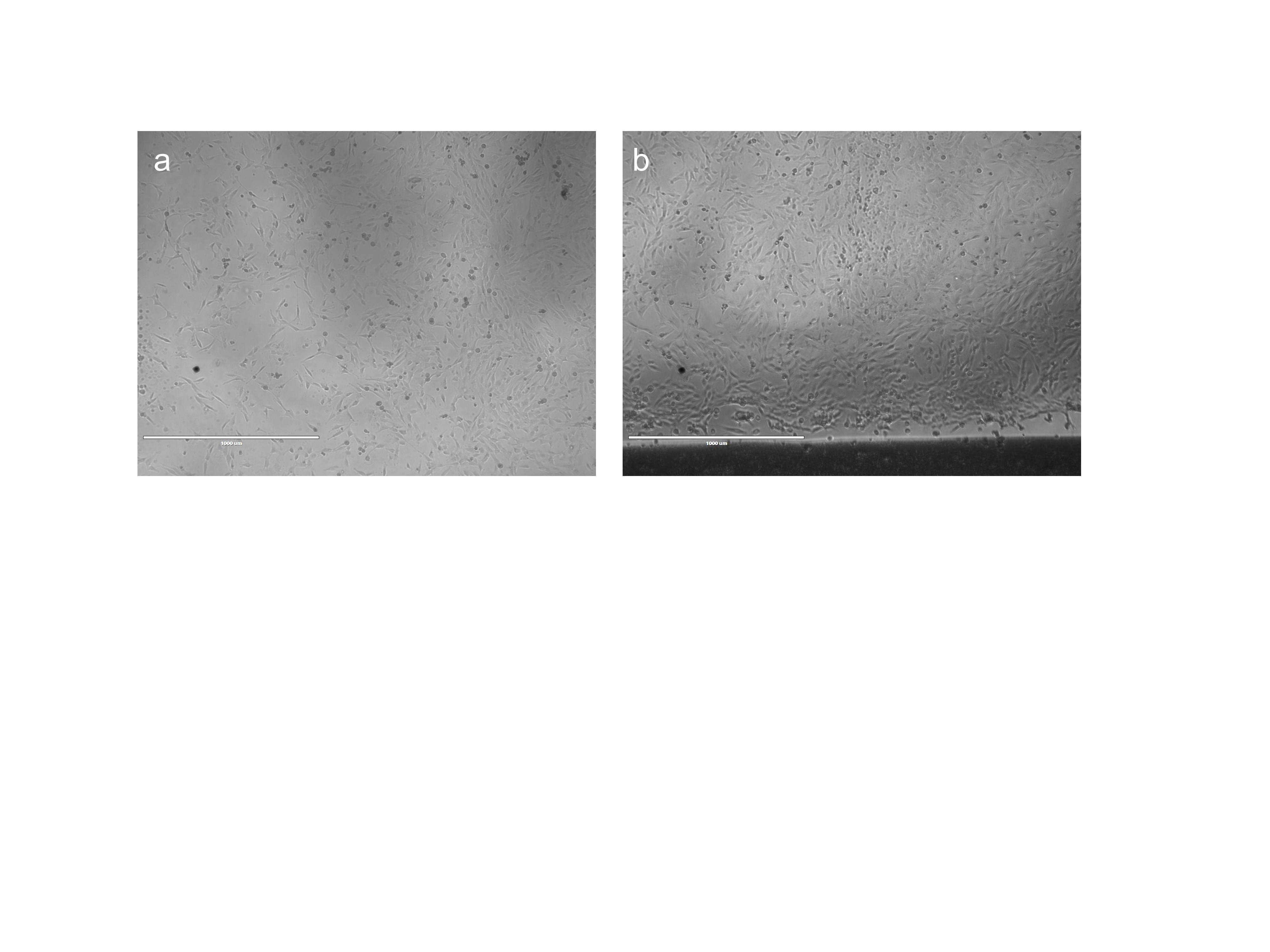
